# Co-expression signatures of combinatorial gene regulation

**DOI:** 10.1101/2020.05.19.104935

**Authors:** Fabio Gomez-Cano, Qian Xu, Shin-Han Shiu, Arjun Krishnan, Erich Grotewold

## Abstract

Gene co-expression analyses provide a powerful tool to determine gene associations. The interaction of transcription factors (TFs) with their target genes is an essential step in gene regulation, yet to what extent TFs-target gene associations are recovered in co-expression studies remains unclear. Using the wealth of data available for Arabidopsis, we show here that protein-DNA interactions are overall poor indicators of TF-target co-expression, yet the inclusion of TF-TF interaction information significantly enhance co-expression signals. These results highlight the impact of combinatorial gene control on such gene association networks. We integrated this information to predict higher-order regulatory complexes, which are difficult to identify experimentally. We demonstrate that genes strongly co-expressed with a TF are also enriched in indirect targets. Our results have significant implications on the empirical understanding of complex gene regulatory networks and transcription factor function, and the significance of co-expression from the perspective of protein-protein and protein-DNA interactions.

## INTRODUCTION

The translation of genotype into phenotype is largely dependent on genes being expressed in the appropriate cell types at the correct time. Such expression is mainly controlled by transcription factors (TFs) recognizing specific *cis*-regulatory regions in the genes that they regulate resulting in protein-DNA interaction (PDI) networks with scale-free properties characteristic of each organism^1^. PDIs are experimentally identified using combinations of gene- and TF-centered approaches; gene-centered approaches result in the identification of TF regulators for specific genes, while TF-centered approaches permit to identify target genes of a particular TF^2–4^. The yeast one-hybrid (Y1H) assay provides the most commonly used gene-centered approach^3^, while TF-centered strategies include chromatin-immunoprecipitation (ChIP) and DNA-affinity purification (DAP) methods, often coupled with high-throughput sequencing (ChIP-Seq and DAP-Seq, respectively)^5,6^.

Identification of PDIs is particularly important in the context of the effect that a TF has on the expression of its target genes. Often, however, identified TF targets show no changes in expression when the activity of the corresponding TF is perturbed^7–11^. While in some instances technical artifacts are responsible, the low overlap between TF targets and differentially expressed genes are more often due to redundancy in the activity of the TF^12,13^, and regulation of the target gene by the TF in only a fraction of the cells sampled. For these reasons, the tethering of a TF to the regulatory region of a gene without a clear contribution to the control of the gene’s expression is often considered of limited biological significance ^14,15^.

Conversely, it is generally assumed that genes with very similar expression patterns are potentially regulated by similar mechanisms, involving shared TFs^16,17^. Similar patterns of gene expression can be captured by gene co-expression networks, in which each node in the network represents a gene, and two nodes can be connected by an edge if they have a significant co-expression relationship^18^. Co-expression relationships can be measured by correlation coefficients, such as Pearson Correlation Coefficients (PCCs), or mutual information (MI) measures, each having advantages and disadvantages^19^. Multiple examples of implementation of co-expression networks or specific TF-target co-expression patterns have allowed the prioritization of specific PDI^14,20^. However, it is unclear to what extent co-expression networks are able to recapitulate PDI networks. Integrating the data of PDIs and gene co-expression networks is not trivial, and researchers usually accept that biologically associated genes and PDIs will show a robust co-expression^21,22^, and/or that genes highly co-expressed are subject to similar regulatory programs^22^. This hypothesis was tested in *Saccharomyces cerevisiae*, where it was shown that two genes have a 50% probability to be controlled by the same TF if the expression correlation coefficient is of 0.84^23^. However, it remains unclear if, in other organisms, co-expression has a similar predictive value to identify TF targets.

TFs function in large complexes, increasing the specificity by which a target gene is recognized, and expanding the regulatory repertoire of a proportionally small number of TFs for a much larger number of genes that they need to regulate. This is generally known as combinatorial gene regulation^24,25^. Components of regulatory complexes assemble through protein-protein interactions (PPIs) and/or PDIs. Models have been developed that use collections of PDIs (or collections of *cis*-regulatory elements) to predict modules of TFs working together^22,26,27^. Similarly, algorithms are available that integrate information from PPIs and PDIs to predict target gene expression, or that combine co-expression with multiple data layers (including PPIs and PDIs) to predict gene regulation^14,28,29^. However, in all these models, the co-expression is used a prioritization tool, and in most instances, the impact of PPIs and PDIs on the expression of the genes that they control must be experimental determined^30^. For the most part, it remains unclear to what extent the co-expression patterns of a TF and its targets is affected by the formation of TF complexes, a general characteristic of regulation of gene expression in eukaryotes^31^.

*Arabidopsis* provides an attractive system to investigate the co-expression relationships between TFs and their experimentally identified target genes in a multi-cellular organism setting. The ATTED-II (http://atted.jp/) database furnishes co-expression information derived from different gene expression analyses^32^. In addition, over five million PDIs have been determined by ChIP-Seq (and ChIP-chip) or DAP-Seq, and most are available through AGRIS (http://agris-knowledgebase.org/)^33,34^. Finally, there are 9,503 experimentally established PPI for *Arabidopsis* TFs that can be accessed through the BioGRID database^35^. Here, we used the wealth of information available in *Arabidopsis* to determine how frequently a TF is co-expressed with its corresponding target genes, and how the co-expression patterns are affected by the formation of TF-TF complexes. We used co-expression analyses at different scales to determine that about half of the TFs are globally co-expressed with their targets as a set, with this number increasing to 85% when local co-expression patterns are considered. We show that a small fraction (in average ~5%) of the direct targets are robustly co-expressed with the corresponding TF. However, when TF complexes deduced from available PPI data are considered, the number of targets co-expressed with a TF significantly increases. By integrating PDIs, PPIs and co-expression information, we predicted the formation of ternary TF complexes, some with strong support from experimental data. Finally, we determined TF’s most highly co-expressed are larger presented by direct and indirect TF targets. Our findings have significant implications on the empirical understanding of complex gene regulatory networks, and the meaning of co-expression from the standpoint of PPIs and PDIs.

## RESULTS

### Transcription factors and their direct targets show varying levels of co-expression

To investigate the co-expression of *Arabidopsis* TFs and their direct target genes, we collected existing PDI data, involving 555 TFs and 25,255 target genes (see *Methods*, Supplementary Table 1). With these dataset, we built a PDI network that included 2,271,066 interactions that were then used to interrogate the co-expression relationships between each TF and its targets, using the mutual rank (MR) of the PCC (MR-PCC), as reported by ATTED-II^32^, and the mutual rank of the mutual information (MR-MI) (See *Methods*). We used PCC and MI capturing linear and non-linear relationships, respectively^36^, and the corresponding MR value in order to reduce dataset-dependent associations and to improve the predictive power of functional associations^32,37^.

To evaluate the significance of the co-expression value for each TF and its entire set of target genes, we carried out two different analyses for each TF: (1) We compared the average MR of a TF with its targets versus the average MR of the TF with a similarly-sized random gene set. Those TFs which showed significant differences with the random set were scored as ‘co-expressed by MR average’ (See *Methods*). (2) We evaluated differences in the distributions of the MRs of a TF with its targets versus all the non-targets genes. Those TF-targets that showed significant differences (*P* < 0.05, Kolmogorov-Smirnov test) from the distribution of the TF-non-target distributions were scored as ‘co-expressed by MR distribution’ (See *Methods*). It should be noted that all the analyses based on MR-PCC values were done independently for negative and positives correlation values. Also, given that we used MR-PCC values and calculated MR-MI co-expression values based on multiple expression experiments (See *Methods*), these results represent TF-target global co-expression patterns. Overall, combining both statistical tests (Supplementary Fig. 11, b), we determined that 231/555 (using MR-PCC) and 172/555 (using MR-MI) corresponded to TFs globally co-expressed with their corresponding sets of targets, with 124 TFs in common between both MR-PCC and MR-MI (Fig. 1a). However, for 276/555 TFs, the co-expression values with their respective targets were not significantly different from what could be expected for a random set of genes, or from non-target genes.

**Figure 1.**
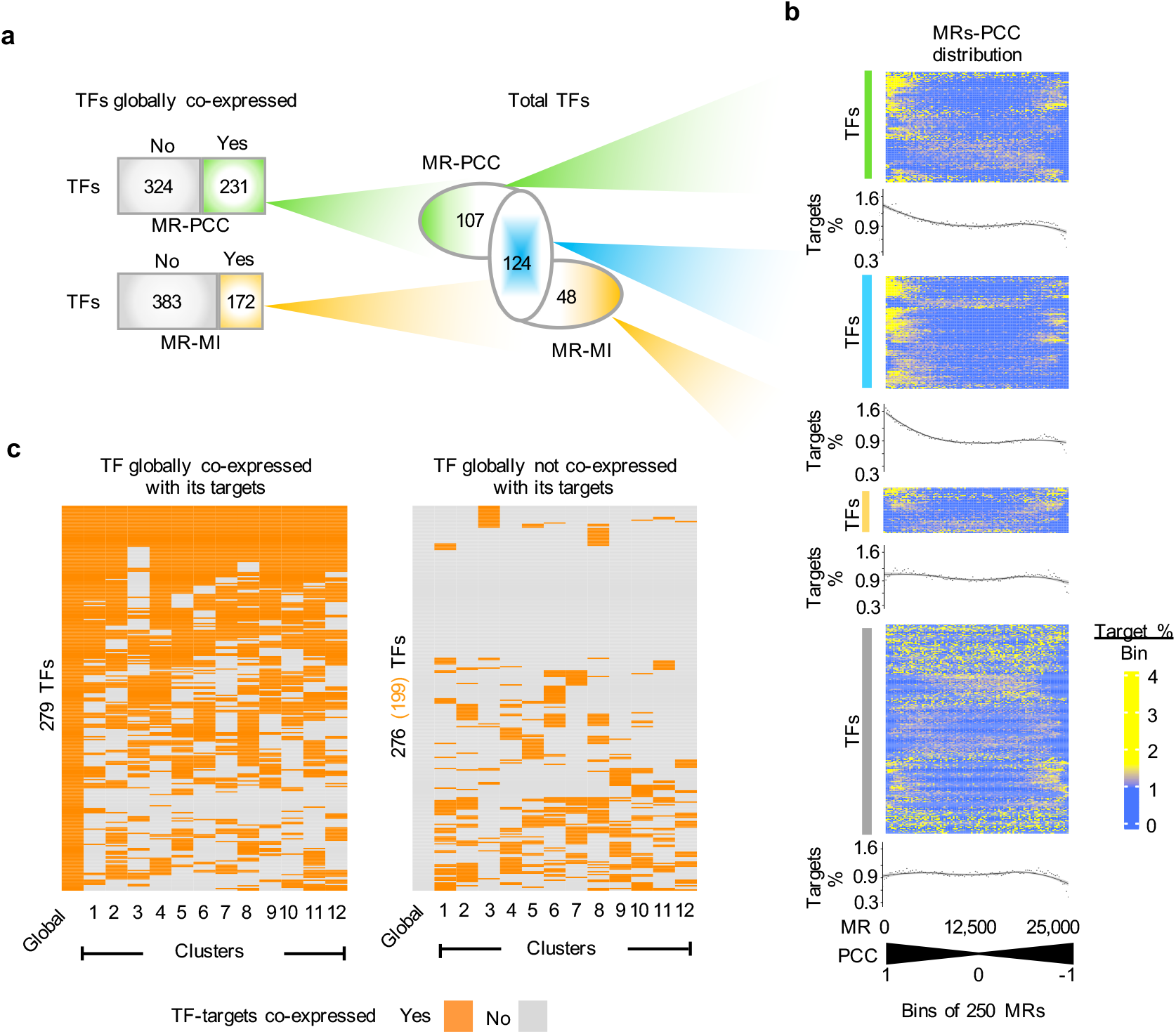
Patterns of co-expression between TFs and their direct target genes. (a) Total number of TFs co-expressed with their corresponding targets across all tissues and conditions (global) based on MR-PCC and MR-MI. The Venn diagrams represent the overlap between both type of metrics. (b) Heat maps displaying the distribution of MR-PCC values across 25,296 *Arabidopsis* genes. TFs were separated into four co-expression groups: TFs co-expressed with its targets based on MR-PCC (107 TFs), on MR-PCC and MR-MI (124 TFs), MR-MI (48 TFs), and TFs that did not shown significant co-expression with its corresponding targets (555 – 107 - 276 – 44 = 276 TFs). Colors represent the percentage of TF targets within bins of 250 MRs. In total, there are 101 bins along the PCC distribution corresponding to co-expression values of each TF with 25,296 genes (genes expressed in dataset used, see *Methods*). Small MR values represent positive PCC values, while large MR values correspond to negative PCC values. Dot plots (seen as curves) under each heat map represent the average percentage of targets for all the TFs for each bin. (c) Heatmap indicating the local co-expression profile of each TFs analyzed along 12 different expression clusters. The color represents binary state of co-expression with yes (co-expressed) in orange and no (no co-expression) in grey. The left panel represents TFs which are co-expressed globally with their targets, while the right panel represents those that are not.

A closer look into the MR-PCC results allowed us to establish that 186/231 TFs showed significant co-expression (either by MR distribution and/or MR average tests) only with positively co-expressed targets (potential transcriptional activators), and 23/231 only with negatively co-expressed targets (potential transcriptional repressors) (Supplementary Fig. 1c). Remarkably, 22 TFs showed significant co-expression with both positively and negatively associated target genes, indicating that they can function both as transcriptional activators or repressors, depending on the target gene subset (Supplementary Fig. 1c).

To further characterize the TFs significantly co-expressed and not co-expressed with their targets (Fig 1a), we evaluated the accumulation of targets genes along the MR distribution based on the PCC and MI metrics. Thus, we first separated the TFs into four co-expression groups: TFs co-expressed with its targets based on MR-PCC (107 TFs), on MR-PCC and MR-MI (124 TFs), MR-MI (48 TFs), and TFs that did not shown significant co-expression with their corresponding targets (276 TFs) (Fig 1a). Then, we binned the MR distribution from the smallest to the largest rank to account for the percentage of total targets genes that fall into each bin. Thus, small and large MR-PCC values corresponded to the most positive and negative co-expression values, respectively. On the MR-PCC distribution, TFs significantly co-expressed with their targets were found to be distributed along the first 24 bins on either the positive or negative values (Fig 1b). This pattern was different for TF-target associations that showed non statistically significant co-expression MR values (Fig 1b, gray panel). Similar patterns were observed for the MR-MI distribution (Supplementary Fig. 2). It was notable that the number of target genes, even in those bins showing the highest absolute vales of co-expression, did not exceed 4-5% of the total targets for any TF, with an ~1% constant number of targets present over all the bins of the distribution (Fig 1b, indicated by the target % curve under every graph). These results indicate that, for *Arabidopsis*, no simple co-expression relationship exists between TFs and their experimentally determined direct targets.

To determine whether signatures of local co-expression could be identified for the 276 TFs that showed no significant global co-expression with their targets, we first grouped the 1,409 gene expression datasets available at ATTED-II into 12 sample clusters (Supplementary Fig. 3), using k-means clustering after a dimensionally reduction of the expression data (See Supplementary Note 1). These 12 cluster were defined as local expression conditions, and by extension were used to re-calculate local MR-PCC and MR-MI values. Using the two statistical tests described above, we determined that 199/276 TFs significantly co-expressed with their corresponding targets in at least one of the clusters (Fig. 1c). As expected, TFs globally co-expressed with their targets were found to be co-expressed in many local clusters (Fig. 1c), with the exception of seven TFs (WIP5, MYB1, PLT1, ERF109, HHO5, NAC4, and AT5G47660), which showed significant global co-expression, but no obvious local co-expression in any of the clusters. The reason for this intriguing behavior is not yet clear.

We investigated what might characterize those TFs that are not co-expressed with their targets globally or locally. We noticed that the connectivity in the network between TFs that show global or local co-expression with their targets compared to those TFs that don’t co-expressed is significantly different. Both the in-degree, which represents how many different TFs bind to a particular promoter region (see *Methods*), as well as the out-degree, which represents the number of target genes bound by a given TF, are significantly smaller for TFs that are not co-expressed with their targets (P < 0.05, Mann-Whitney U test; Supplementary Fig. 4). While these results may suggest that lowly-connected TFs in the network exhibit a different co-expression relationship with their targets, we cannot yet rule out that the clusters might not be ‘local enough’ for these TFs.

### Few targets are highly co-expressed with their respective TFs

The accumulation of target genes along the MR-PCC distribution previously described (Fig. 1b) indicates a low abundance of targets among the genes most highly co-expressed with each respective TF. Indeed, the maximum percentage of targets within a bin of 250 co-expressed genes is ~5% (Fig. 1b), and the percentage of targets by bin decreases after the first 5,000 MRs, which captures at most 25% of the total direct targets identified for each TF.

To evaluate the percentage of highly co-expressed targets (HCT) for each TF, we defined the top and bottom 2.5% of the MR-PCC distribution as the set of highly co-expressed genes (HCGs), and then we counted the total number of targets in these intervals. The TF that has the maximum percentage of target genes (36%) identified as HCTs using the above criteria was found to be ARABIDOPSIS PSEUDO-RESPONSE REGULATOR 9 (PRR9). However, on average, only 4.7% of the targets corresponded to HCTs (Fig. 2a), indicating that (on average) the remaining 95.3% of the targets corresponded to low co-expressed targets (LCTs).

**Figure 2.**
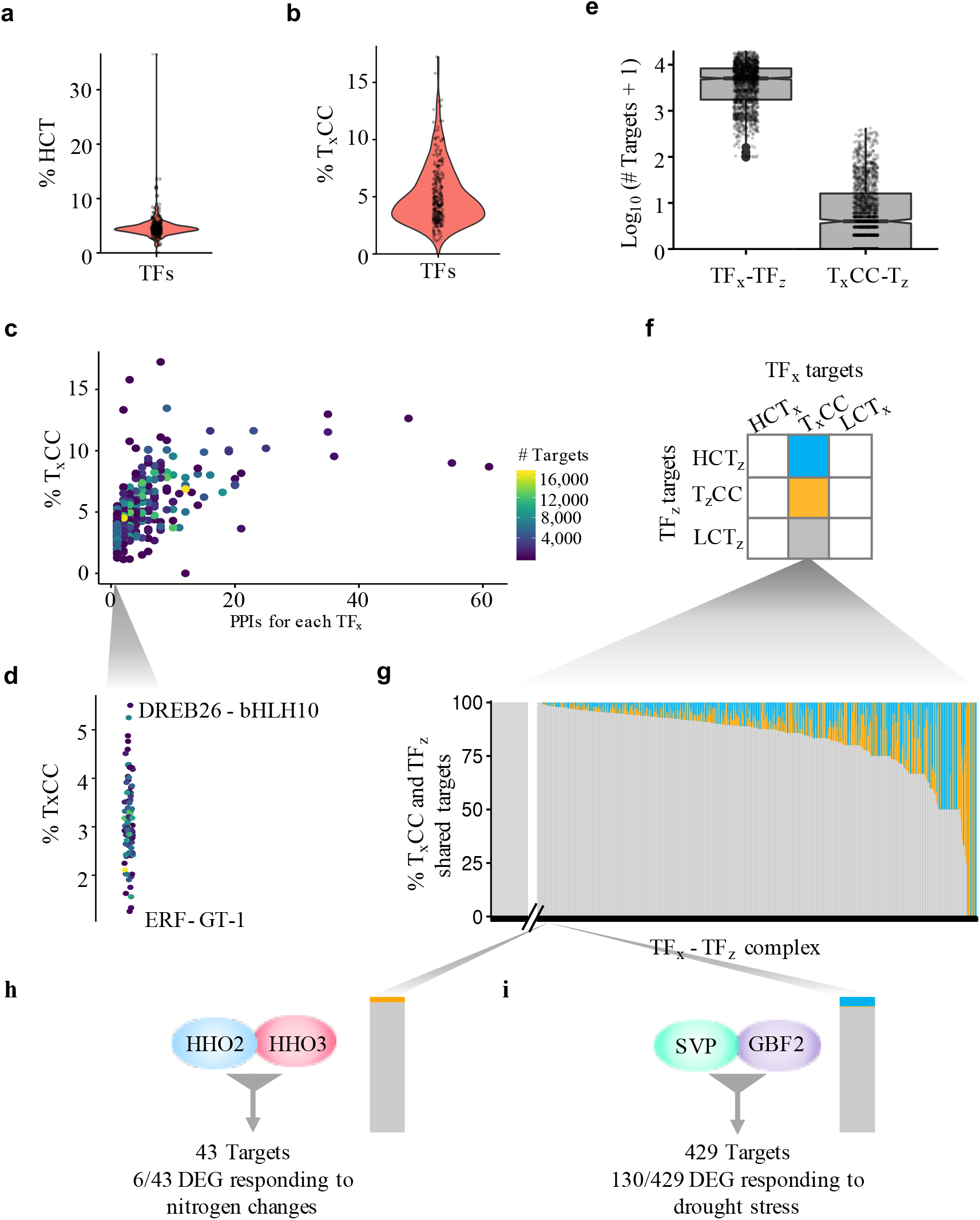
Targets are more frequently co-expressed with TF complexes, but not with individual TFs. **(a)** Violin plot showing the percentage of highly co-expressed targets (HCT) for 313 TFs. **(b)** Boxplot representing the percentage of low co-expressed targets (LCTs) that correspond to targets co-expressed with a TF_x_ complex (T_x_CC). **(c)** Percentage of T_x_CCs as a function of the total number of PPIs in which each TF is involved. **(d)** Magnification of the section in (c) corresponding to TFs with just one interacting partner. DREB26-bHLH10 and ERF (At4g18450)-GT-1 represent the extreme upper and lower cases on the distribution. The color scale in (c) and (d) represent the number of targets for each TF. **(e)** Boxplot showing the number of shared targets between the 815 TF complexes analyzed and the number of targets of a given TF_x_ co-expressed with the complex TF_x_-TF_z_ (T_x_CC) that are also targets of TF_z_. **(f)** Venn diagrams displaying how targets of a TF_z_ are distributed among the HCTs, TCCs, or LCTs of a TF_x_ represented in blue, orange and yellow, respectively. **(g)** Distribution of targets according to the comparison in (f) for the of 815 TF_x_-TF_y_ complexes analyzed. Complexes are along the x-axis, while the y-axis indicates the overlapping frequency. The HHO2-HHO3 **(h)** and SVP-GBF2 **(i)** TF complexes provide representative examples from the 815 TF complexes analyzed. **(h)** The numbers indicate the differential expressed genes (DEG) under different nitrogen growth conditions. **(i)** The numbers indicate DEGs, also predicted as targets of the corresponding complexes, under drought stress in three different studies. Side bar plot represents a zoom over the HHO2-HHO3 and SVP-GBF2 position on the shared target distribution displayed in **(g)**.

### PPIs condition TF co-expression with direct targets

To better understand why so few targets are highly co-expressed with any given TF, we investigated how having multiple, often physically interacting, TFs controlling any given gene might influence binary TF-target co-expression metrics. For this, we collected from BioGRID 815 experimentally determined PPIs involving 313/555 TFs studied here. Using this PPI information, we evaluated to what extent the formation of a TF complex (*e.g.,* TF_x_-TF_z_) explains the large fraction of LCTs for each TF_x_by calculating the partial co-expression correlation of TF_x_ with all the LCTs, conditioned on TF_z_^38–40^. It should be noted that such correlations are not symmetric, meaning that T_x_CC could be different from T_Z_CC. We assessed the correlation with all the *Arabidopsis* genes to define the top 2.5% (at each tail of the correlation distribution) of highly co-expressed genes for each TF_x_-TF_z_ complex. We found that, in average, 5% of the LCTs of a TF are co-expressed with the complexes in which the TF is involved (Fig. 2b), and the range of co-expression of the LCTs with TF complexes (0% - 17.2%, Fig. 2b) does not depend on the total number of targets of the corresponding TF (Spearman Correlation, *r_s_* = 0.02) (Fig. 2c, color scale distribution). Notably, the percentage of targets co-expressed with a complex increased as a function of the number of interactors that a TF has (correlation, *r_s_* = 0.69) (Fig. 2c), indicating that a significant proportion of the LCTs described previously can be explained by considering complexes of interacting TFs.

Even among TFs known to form similar number of complexes, there is significant variation in the proportion of LCTs that co-express with the complex (Fig. 2d). For example, when we focused on TFs that have just one known partner, we observed that the extreme examples in this group corresponded to DEHYDRATION RESPONSE ELEMENT-BINDING PROTEIN 26 (DREB26) and RELATED TO AP2 11 (RAP2.11), which interact with BASIC HELIX LOOP HELIX PROTEIN 10 (BHLH010) and ELONGATED HYPOCOTYL 5 (HY5), and for which the complex explain 11.3% and 2.7% of LCTs, respectively (Fig. 2d). This result shows that each TF complex has a specific and unique effect on the percentage of targets that are co-expressed, possibly reflecting functional aspects of combinatorial gene regulation.

So far, we have shown that the co-expression of TFs with their targets can be enhanced by considering the regulatory complexes in which each TF is involved. Although, all these interacting TFs share a variable number of common targets (TF_x_-TF_z_ in Fig. 2e), only a small fraction of the shared targets are co-expressed with the complex (T_x_CC-T_z_, Fig. 2e). Thus, to better understand the co-expression behavior of the TF_x_ and TF_z_ shared targets, we compared which fraction of the targets common to both TF_x_ and TF_z_ that co-express with the TF_x_-TF_z_ complex are also highly co-expressed with TF_z_ (Fig. 2f, blue box), are co-expressed with the complex TF_z_-TF_x_ (Fig. 2f, orange box), or are among the low co-expressed targets with TF_z_ (Fig. 2f, grey box). Overall, 91% of the shared targets of TF_x_ and TF_z_ that are co-expressed with the TF_x_-TF_z_ complex (T_x_CC) correspond to targets of TF_z_ that show modest co-expression with TF_z_ (LCT_z_, gray in Fig. 2g). Only 3.9% of the TF_x_-TF_z_ shared targets is highly co-expressed with TF_z_ (Fig. 2g, blue box), and 4.1% with both complex (T_x_CC and T_z_CC) (orange in Fig. 2g). These results underscore the importance of considering TF complexes when interpreting the (lack of) co-expression between TFs and their targets.

To evaluate the biological relevance of the observed co-expression of targets with TF complexes, but not with the individual TFs, we focused our attention on a few examples. HHO2 (HRS1 HOMOLOG2) and HHO3 (HRS1 HOMOLOG3) encode MYB-related TFs involved in phosphate homeostasis and lateral root development^41^, which also participate in nitrogen responses^42^. Our analysis showed that the HHO2-HHO3 complex co-expressed with 43 targets. Note, HHO2 and HHO3 as well as six of its targets are differentially expressed genes that respond to different N growth conditions (Fig. 2h), supporting the idea of functional relevance of complex formation and its targets. We also analyzed the SHORT VEGETATIVE PHASE (SVP) - G-BOX BINDING FACTOR 2 (GBF2) complex. SVP is a flowering repressor^43^ related also to drought responses^44^. GBF2 has been related to abscisic acid (ABA) responses^45^. Our results indicated that the SVP-GBF2 allow us to identify 429 shared co-expressed targets (Fig. 2i), of which 130 genes were also differentially expressed under drought responses^44,46,47^. Altogether, these results provide further evidence that an important fraction of TF targets that do not significantly co-express with the TF are indeed co-expressed when TF complexes are considered.

### Co-expressed targets shared by binary TF complexes suggest higher-order arrangements

Our results indicate that the integration of co-expression and physical interaction information contributes to the identification of TFs that control gene expression working as part of complexes. There are many instances in which *Arabidopsis* TF pairs interact and control shared sets of target genes^25,48^. However, the experimental identification of higher-order TF complexes is not without challenges^49^.

To investigate whether the combination of co-expression, PPI, and PDI information might provide insights on higher-order TF complexes, we started by describing the complexes made up by TGA10 (TGACG MOTIF-BINDING PROTEIN 10), TCP14 (TGA10 with TEOSINTE BRANCHED, cycloidea and PCF 14), and a homeodomain-like TF (AT2G40260)^50^. The TGA10-TCP14 and TGA10-AT2G40260 complexes share 80% of targets co-expressed with each complex (Fig. 3a, black nodes). Moreover, shared targets had similar expression correlation with both heterodimers (either positive or negative), indicating that both complexes potentially activate or repress the same sets of genes (Fig. 3a). These results, combined with the information that TCP14 and AT2G40260 physically interact with each other^50^, provide strong evidence that TGA10, TCP14, and AT2G40260 form a ternary complex that controls the expression of all targets indicated in Fig. 3a.

**Figure 3.**
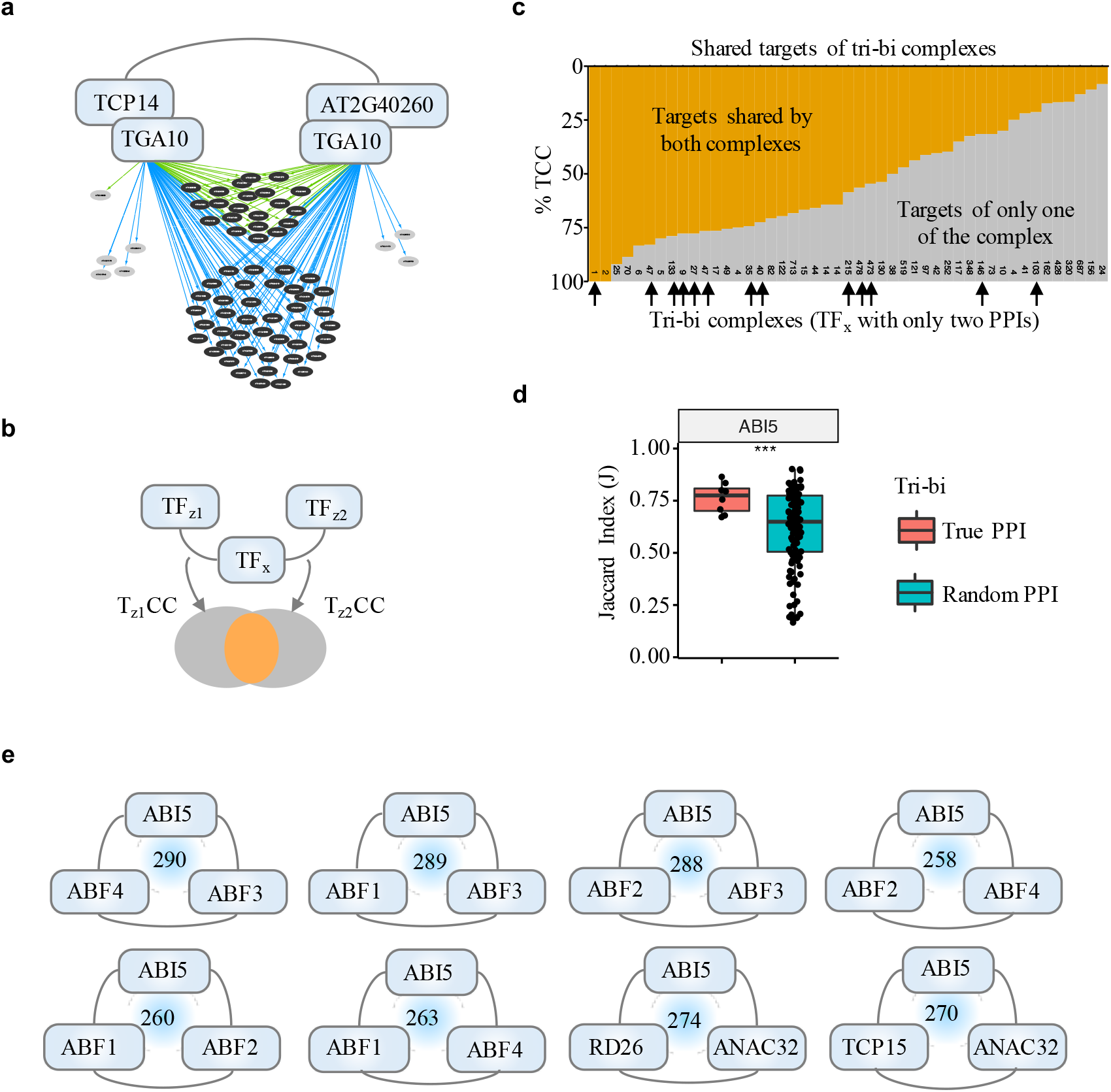
Common co-expressed targets of TF complexes suggest higher-order TF arrangements. **(a)** Shared co-expressed targets for the TGA10-TCP14 and TGA10-AT2G40260 TF complexes. Common targets for both complexes in each instance are indicated in black, while those controlled by one complex, but not by the other are in light gray. Green arrows show targets with a positive co-expression correlation (indicative of activation), while those in blue correspond to targets with a negative co-expression correlation (indicative of repression) with the respective TF complexes. (**b**) Schematic representation of the strategy used to identify shared targets by comparing T_x_CC between pairs of dimers. **(c)** Percentage of the total targets that are bound by both complexes (orange) or just by one (gray). Black arrows indicate tri-bi complexes with PPI information for all three binary interactions. **(d)** ABI5 as one example of 12 TFs for which the fraction of shared targets of the tri-bi complexes are significantly larger (two-sided *t*-test *P* < 0.05) than tri-bi complexes formed by random interactions. Similarity between the sets of target genes of corresponding dimers were measured by computing the respective Jaccard indices. **(e)** Tri-bi complexes involving ABI5. Experimentally-verified interactions are indicated by lines, and targets of the complexes are indicated by the numbers in blue.

We next investigated how many other instances of such ternary pairs (tri-bi, from here on, for triple-binary) of TFs might be present in *Arabidopsis*. For this, we first identified 47 TFs with at least two interactors and with PDI information to determine the percentage of shared target genes between both pairs (Fig. 3b, orange), and compared the total percentage of shared target with the percentage targets of just one pair (Fig. 3b, gray). In some instances, all targets are shared by the two binary complexes (all orange columns, Fig. 3c), while only ~8% are shared by those binary complexes with the smallest overlap (columns to the right, Fig. 3c). Providing supporting evidence for the formation of higher-order (ternary) complexes, for 13/47 tri-bi tested there is PPI information for all three binary interactions (indicated by black arrows in Fig. 3c). However, we could not establish a statistically significant correlation linking the number of shared targets and experimental evidence confirming the formation of ternary complexes. While it is possible that the percentage of shared co-expressed targets is not a good indicator of the formation of ternary complexes, the lack of a statistically significant correlation more likely reflects sparse PPI data for many of the TF pairs involved.

We next investigated how frequently TFs involved in tri-bi interactions share common targets. In total, we found 2,013 true tri-bi instances *(i.e.,* with evidence of physical interaction for all pairs of the tri-bi) involving 140 TFs. In ~90% of these cases, the TFs involved showed a significant overlap of target genes (false discovery rate < 0.01, Fisher’s exact test) (Supplemental Table 2), indicating that TFs involved in tri-bi interactions which share a significant fraction of targets are excellent candidates for the formation of tertiary, or even higher-order, complexes. To determine if the fraction of shared targets differ from random tribi complexes, we compared the shared targets co-expressed of TF complexes from tri-bi instances experimentally demonstrated PPI versus tri-bi instances obtained from randomized binary interactome for each TF (see *Methods*).

Out of the 104 TFs, we found 12 TFs involved in tri-bi instances with a fraction of shared targets that was significantly larger (see *Methods*) than expected from the background model (Supplementary Fig. 5a). An example is provided by ABI5 (ABA INSENSITIVE5, At2g36270), which is involved in eight tri-bi instances with a median shared fraction of targets of 0.77 (Fig. 3d). Notably, six out of the eight tri-bi instances involving ABI5 were composed by a combination of four TFs of the same family (ABF, ABSCISIC ACID RESPONSIVE ELEMENTS-BINDING PROTEIN) (Fig. 3e). The number of target genes for each tri-bi instance ranges from 258 for ABF2-ABI5-ABF4 to 290 for ABF3-ABI5-ABF4 (Fig. 3e). The 290 ABF3-ABI5-ABF4 gene targets include 46 genes differentially expressed in *abi5* mutant seeds^51^. Remarkably, ABF2, ABI5, and ABF4 also interact with SnRK2.2 (SNF1-RELATED PROTEIN KINASE 2), PP2CA (PROTEIN PHOSPHATASE 2CA)^52,53^, and AHG1 (ABA-HYPERSENSITIVE GERMINATION 1)^52^, which are key known posttranslational regulators of ABI5^54^. We also found 41 TFs involved in tri-bi instances with a fraction of shared targets that were significantly lower than expected from the background model (Supplementary Fig. 5b). Assuming that the binary PPIs that form these bi-tri instances occur *in vivo*, these results strongly suggest that they involve TFs that bind overlapping sets of target genes as part of dimeric complexes.

### Genes highly co-expressed with TFs are enriched in indirect TF targets

In previous sections, we focused on the patterns of co-expression between TFs and their direct targets. However, a question that remains unanswered is whether there is a relationship between a TF and the genes that are most highly co-expressed with that TF. To address this, we determined how many target genes of a TF are also among the top 5% most highly co-expressed gene (HCG) with the TF in question. Strikingly, for 80% of the TFs, less than 30% of the HCG are among the target genes, although exceptionally [for NF-BY2 (nuclear factor Y, subunit B2)]^45^ this number can be as high as 82% (14.3% in overall average) (Fig. 4a). We explored the possibility that genes that are not direct targets of a TF_x_ could be targets of a TF_x_ partner (TF_z_), or that they could be targets of a second TF (TF_y_) that is itself direct target of TF_x_.

**Figure 4.**
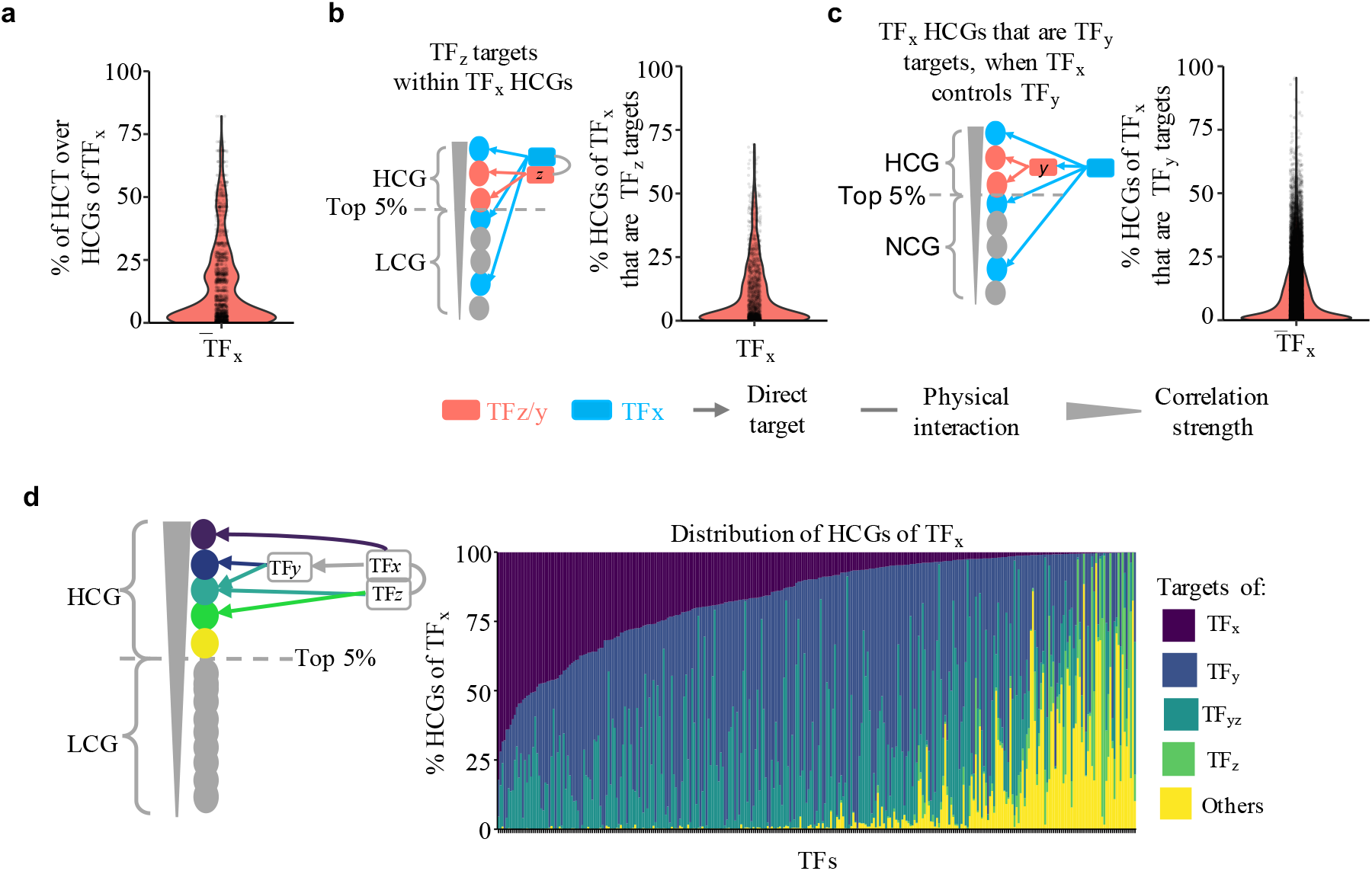
Genes highly co-expressed with TFs are enriched in indirect TF targets. **(a)** Percentage of highly co-expressed genes (HCGs) of TF_x_ that are actual targets of TF_x_. **(b)** Model and percentage of highly co-expressed genes that are potential indirect targets of TF_x_ because of indirect action of a TF_z_ interactor of TF_x_. **(c)** Model and percentage of highly co-expressed genes that are targets of a TF_y_ regulated by TF_x_. **(d)** Percentage of HCGs explained as direct or indirect targets of TF_x_.

To determine the contribution of the TF partners (TF_z_) to the set of genes highly co-expressed with TF_x_, we evaluated how many of highly co-expressed genes of TF_x_ are targets of any TF_z_, but not of TF_x_ itself. In total, we found that 309 TFs (out of the 313 TFs tested) had at least one highly co-expressed gene that was target of the any of its TF_z_ partners. On average, we determined that 10% of the highly co-expressed genes of a TF correspond to this class (Fig. 4b) (Supplementary Table 4).

To establish the contribution of indirect targets to the set of genes most highly co-expressed with a TF in regulatory hierarchy (TF_y_), we analyzed the same set of 313 TFs. From these TFs, 306 bind a TF_y_ which had at least one direct target also highly co-expressed with the upstream TF_x_. On average, 9.8% of the genes most highly co-expressed with a TF_x_ corresponded to indirect targets of TF_y_ (Fig. 4c). We compared the actual set of HCGs recovered based on true interaction versus random networks (of PPI and PDI, respectively) (See *Methods*). Overall, random TF PPIs recover a similar number of HCG than known PPIs (P < 0.05, Mann-Whitney U test) (Supplementary Fig. 6a). Note, the PPI network used here does have an average path length of 3.5 edges between all its nodes (TFs), which is an indication of a weak independence between the true and the random PPIs. In contrast, random target TF_y_ recovered a significantly smaller number than the ones recovered by true targets (P < 0.05, Mann-Whitney U test) (Supplementary Fig. 6b), suggesting that indirect hierarchical regulation plays a major role in the explanation of HCG as indirect targets.

We calculated the total contribution of direct and indirect targets to the set of highly co-expressed genes for each of 313 TFs by adding up the contribution of all its interactors (TF_z_) and downstream TFs (TF_y_) (Fig. 4d). Taken together, our results indicate that on average ~90% of the genes most highly co-expressed with a TF are direct targets of the TF (~16%), direct targets of a partner of the TF (~4%, after removing partners which are also direct targets of the TF_x_), and indirect targets (~70%, targets of a TF target) (Fig. 4d). Interestingly, in a 26% from the total, the partner for TF_x_ is also a downstream target, participating in a of feed-forward loop (FFL). FFLs are among the most highly represented regulatory motifs present in *Arabidopsis*^43^ and other eukaryotes^55^.

## DISCUSSION

Our study investigated complex co-expression relationships between TFs and their targets, taking advantage of the extensive PDI, PPI and gene expression data available for *Arabidopsis*. We show that approximately 50% (279) of the 555 TFs investigated are globally co-expressed with their targets as a set, while an additional 35% (199) are co-expressed with their targets in at least one of the 12 clusters into which we grouped the ~1,400 RNA-Seq experiments available. This means that for 77 *Arabidopsis* TFs for which there is extensive PDI information, there is no evidence that they are co-expressed with the identified targets any better than with random sets of genes. Given that many TFs did not show global co-expression with their targets, but only when specific subsets of the expression data was used, the possibility exists that using single cell sequencing may reveal co-expression relationships that are masked by the complexity of the cell populations used in organ-level gene expression experiments.

We show that only a small fraction (< 36%, in average 4.7%; Fig. 2a) of the direct targets are among the genes most highly co-expressed with a given TF. Conversely, direct targets are a small fraction of the genes highly co-expressed with a TF (<82%, in average 14.3%; Fig. 5a). Given that high co-expression is often used as an additional criterium to assert the biological significance of a PDI, our results indicate that these comparisons should be used with significantly more caution.

In an effort to determine the co-expression relationships between TFs and their targets, we noticed that up to 17% of the not so highly co-expressed targets of one TF are in fact co-expressed targets of TF complexes. Indeed, a large portion (up to 100%, in average ~22%) of the targets co-expressed with a TF complex were common targets of members of the complex, which are not highly co-expressed with individual TFs. These findings are consistent with the ample literature describing gene combinatorial control^25,30,56,57^. We explored the biological relevance of the targets co-expressed with two different TF complexes, HHO2-HHO3 and SVP-GBF2, by investigating their expression changes under stress conditions. Remarkable, in both cases we identified target genes and TF members of the complex that were differentially expressed. While intuitively logical, our results provide firm evidence that, to fully exploit co-expression analyses, the combinatorial nature of gene regulation needs to be considered.

Experimentally identifying ternary TF complexes is far from trivial. Using co-expression information combined with PPIs and shared targets derived from PDI data, we carried out an analysis of possible TF pairs that could be part of ternary TF complexes (Fig. 4c). As an example, we identified eight potential ABI5 ternary complexes that involve four out of seven TFs from the same family (ABF1/2/3/4). These results are consistent with experimental data suggesting redundant functions of ABF3 and ABI5^58^, and the regulatory role of ABI5 and ABF2/3/4 on the degradation of chlorophyll related genes^59^. In addition, it is known that ABF3/4 and NF-YC (nuclear factor Y subunit C) form a complex that is able to control flowering in response to drought by regulating expression of SOC1 (SUPPRESSOR OF OVEREXPRESSION OF CONSTANS1)^60^, which is also a target of ABI5 in seedlings^6^. These results strongly suggest the formation of a larger-order complex involving ABF3-ABF4-ABI5. Together, by integrating PPIs between TFs with co-expression studies, we were able to predict a number of potential ternary TF complexes, which could now be experimentally validated, an easier undertaking than carrying out *do novo* identification.

Another question addressed by this study regards the nature of the association of the other genes that are highly co-expressed with a TF, if they are not targets of the TF itself. We showed that, in average for the 313 TFs investigated, almost a third of the highly co-expressed genes, are either indirect targets of the TF (targets of a TF target), direct targets of the TF or direct targets of a TF partner. Is important to note that in many instances this number was much larger, which to some extent justifies the wide-spread use of co-expression as a proxy to carry out functional association of TFs and different plant traits^29,61^. However, what our studies also show is that the use of co-expression is a poor indicator of direct interactions between TFs and their target genes. Establishing the co-expression relationships of TFs and their target genes has wide implication for elucidating the architecture of gene regulatory networks in all organisms, and establishing the meaning of co-expression as a tool to elucidate molecular interactions.

## METHODS

### Data collection

Expression and global co-expression data were collected from the ATTED-II database (http://atted.jp/, versions Ath-r.v15-08 and Ath-r.c2-0, respectively)^32^. In total, we used 1,416 different RNA-Seq libraries with expression data associated for 25,296 different genes. We collected the protein-DNA interaction information as raw peaks (bed or narrowpeak files from ChIP-chip, ChIP-Seq, and DAP-Seq experiments) from the Gene Expression Omnibus (GEO) and/or supplementary material from reference source (Supplementary Table 1). The assignment of a peak region to a gene was carried out assuming a promoter region of 2 kb upstream from the transcription start site (TSS) for each *Arabidopsis* gene (genome annotation TAIR10). We used all peak region sizes as reported originally. All protein-protein interactions (PPIs) used for the identification of complex co-expressed targets were collected from the BioGRID database for Arabidopsis (V3.5.169)^35^.

### Evaluation of co-expression and determination of mutual rank values

For the evaluation of the global co-expression between TFs and their corresponding targets, we used the mutual ranks (MRs) of the Pearson Correlation Coefficient (PCC) and the Mutual information (MI) as co-expression metrics. Both metrics were represented as the MR of the correlation value between compared genes as follows:

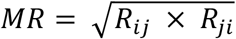

where R_ij_ is the rank of the correlation of gene *i* with the gene *j*, and R_ij_ is the rank of the correlation of gene *j* with the gene *i*, with the highest value as the best rank (close to 1). Global MRs from positive PCC were used as reported by ATTED-II, while global MRs from negative PCC values were transformed into a second MR by subtracting the original MR reported from the maximum possible MR (25,296) for each TF. For the calculation of local MRs-PCC, we used the expression normalized as reported by ATTED-II, parsing the samples into twelve expression conditions through a dimensional reduction of the total dataset, followed by a k-means analysis (see Supplementary Note 1). Grouping these samples as expression conditions, we proceeded to calculate the PCC between genes. This correlation was weighting the samples similarities based on the correlation between samples to avoid an overestimation of the correlation between genes guided by replicates. We calculated the weighted PCC using the R package wCorr (Version 1.9.1)^62^, with an optimal threshold of 0.4, as used by ATTED-II on the global MR-PCCs. All global and local co-expression analyses using MR-MI values were carried out with the same samples used for the calculation of the respective MR-PCC values. The correlation-based on MI was estimated using the R package Parmigene (Version 1.0.2)^63^, and with 1e-12 as noise to break ties due to limited numerical precision.

### Identification of TFs co-expressed with the corresponding target genes

The significance of the MRs between TFs and their corresponding targets was assayed using both MR-PCC and MR-MI correlation metrics, and two independent statistics tests: (1) We compared for each TF the average MR value of the targets vs. a null distribution of average MRs values from 1,000 random sets of genes, referred as co-expression by MR average. Each random sample was generated by sampling with replacement *N* random genes to the *N* number of direct targets of each TF. For the MR-PCC values, we compared separately MR distributions of positively and negatively PCC values. To define if average MRs of the target genes were significantly smaller than the null distribution, we calculated the Z-score using the MR values of the true targets using the random set of genes as background (which follow a gaussian distribution) to then ask if the P value of the Z score was significant (*FDR* < 0.05 after correcting for multiple testing by Benjamini-Hochberg method)^64^. (2) We evaluated the differences between target and non-target genes by testing if their empirical cumulative distribution was similar (*FDR* < 0.05, one-sided Kolmogorov-mirnov test, alternative: greater), referred to as co-expressed based on MRs distribution. In both casas (co-expression base on average and distribution, we did test positive and negative correlation independently).

### Identification of targets co-expressed with TF complexes

The identification of complex-co-expressed targets was carried out for TFs present in our list of TFs with PDI data and at least one protein-protein interaction (PPI) between them in BioGRID. In total, we found 815 protein-protein interactions (PPIs) associated with 313 different TFs. Using these PPIs, we evaluated the effect of the formation of a TF complex (TF_x_-TF_z_) over lowly co-expressed targets (LCTs) of TF_x_ by: (1) Assuming TF_x_-TF_z_ as a new protein, thus, we averaged their expression (TF_x_ and TF_z_) and then re-calculated the co-expression of the complex with a target *y*. This co-expression analysis was carried out using the weighted PCC as described above. (2) We also calculated the partial correlation of TF_x_ with genes *y* conditioned by TF_z_: p(TF_*x*_ ~ y | TF_z_), such that TFx and TF_z_ interact between them and y is a TFx target. The partial correlation was calculated using the R package PPCOR^40^. In both cases, we calculated the co-expression of the complex against all genes in the genome to define the significant values on the distribution obtained (See below).

### Definition of highly co-expressed targets

We defined highly co-expressed genes as those genes in the top 5% of the correlation distribution, assuming them as genes with correlation values significantly different from the average of correlation distribution (*P* < 0.05). For PCC values, we took the 2.5% from each tail, while for MI values we took the top 5%. The approach was also implemented to define highly target co-expressed with a complex (TCC).

### Degree network connectivity

We defined the in-degree and the out-degree as the number of TFs that bound the promoter of a particular target genes and the number of targets of a particular TF, respectively. Differences in both degrees, in-& out-degree, between TF co-expressed with its corresponding targets and those than not were tested by a Mann-Whitney test.

### Protein-Protein Interactions (PPIs) and Protein-DNA interactions (PDIs) network randomization

We created random PPIs and PDI networks to test the significance of the shared targets between dimers of the tri-bi and to test the significance of number the indirect targets within the set if genes highly co-expressed with a TFs, as well as significance of number the indirect targets by TFs in cascade. In all the cases we used the *rewire* function from the R package Igraph (v1.2.4.1) to generate the random network with similar degree by node and avoiding loops (niter=NodesInNetwork*1000). Random PPI network was built with *directed* set as *FALSE* while the random PDI was set as *TRUE*, which allow the shuffling of edges between TF and target genes only.

### Definition of tri-bi complexes with significant number of shared targets

In total, we selected 104 TFs after discarding tri-bi instances with no significant target overlap, as well as TFs involved in less than two tri-bi instances (to avoid comparison with few samples). To compute the differences between the random and true PPIs, we calculated the Jaccard index (J) between every pair of dimers involved in each tri-bi, and then we asked if the mean of the J values between true tri-bi instances was different from the J values mean of tri-bi instances derived from the random PPI collection (see randomization network description).

### Definition of significant number of indirect of TF_z_ and TF_y_ within HCG of a TF_x_

To test the significance of the percentages of HCG of TF_x_ explained because either they are targets of an interactor TF_z_ or a target TF_y_; we compared the actual set of HCGs recovered based on true interaction versus random networks (of PPI and PDI, respectively). We measured the overlap (Jaccard index) of the HCGs of TF_x_ with the corresponding set of TF_z_ and TF_y_ targets.

## ACKNOWLEDGEMENTS

This work was supported in part by grants to E.G. from the NSF IOS-1733633, the DOE DE-FOA-0001650, to S.-H.S. from NSF IOS-1546617, DEB-1655386, and DOE Great Lakes Bioenergy Research Center (BER DE-SC0018409), and to F.G.C. from MSU and NSF Research Traineeship Program (DGE-1828149).

## AUTHOR CONTRIBUTIONS

F.G.-C. carried out the majority of the analyses with the assistance in the statistical analyses by Q.X. and A.K.; F.G.-C. and E.G. wrote the manuscript with contributions from all the authors and S.-H.S. contributed with critical discussions; E.G. coordinated and supervised the research project and agrees to serve as the author responsible for contact and communication.

## COMPETING INTERESTS

The authors declare no competing interests.

## SUPPLEMENTARY FIGURE LEGENDS

**Supplementary Fig. 1.** Evaluation of co-expression of TFs and corresponding target genes as a set. Comparison of the two statistical approaches used to test differences in either average or distribution of MRs between targets and not targets genes by **(a)** PCC-MR or **(b)** MI-MR. **(c)** Venn diagrams comparing the total number of positive and negatively co-expressed TFs with their targets based on PCC-MR.

**Supplementary Fig. 2.** Heat maps displaying the distribution of MR-MI values across 25,296 *Arabidopsis* genes. Colors represent the percentage of TF targets within bins of 250 MRs. In total, there are 101 bins along the PCC distribution corresponding to co-expression values of each TF with 25,296 genes (genes expressed in dataset used, see *Methods*). Small MR represent largest MI, thus, better association between TF and genes in bin. Dot plot under each heat map represent the average percentage of targets for all the TFs along each bin. Color side bar represent TFs categories as presented in Fig 1.

**Supplementary Fig. 3.** Sample expression clusters used to define local expression values. Clusters were defined by k-means clustering (k=12 define by Elbow method) using the t-Distributed Stochastic Neighbor Embedding (t-SNE) 1 and 2 of the expression data (See Supplemental Note 1).

**Supplementary Fig. 4.** In- and out-degree differences between TFs co-expressed and not-co-expressed with their targets. This classification accounts for both globally and locally co-expression results. Both type of degree (in and out) showed statistically significant differences between TFs co-expressed or not co-expressed with its targets (Mann-Whitney U test, P. value < 0.05).

**Supplementary Fig. 5.** Comparison of target genes recovered for tri-bi of 53 TFs with a shared fraction significantly larger **(a)** or smaller **(b)** than by random PPIs. The similarity of the recovery set of targets was measured as the Jaccard index between the set of targets of each pair of dimers that form a tri-bi complex. Asterisks indicate P-value significance (*: p <= 0.05, **: p <= 0.01, ***: p <= 0.001, ****: p <= 0.0001, two-sided *t*-test).

**Supplementary Fig. 6.** Comparison of HCG which are not targets of TF_x_ recovered because they are either **(a)** a target of a TF_z_ interactor of TF_x_, or **(b)** a target of a TF_y_ regulated by TF_x_ vs random interaction. Jaccard index (J) calculated as the number of TF_z_/TF_y_ targets shared with the HCGs non-targets of TFx over the total TFz/y targets plus total HCGs no-targets.

## LEGENDS OF SUPPLEMENTARY TABLES

**Supplementary Table 1.** List of TFs analyzed in this study.

**Supplementary Table 2.** Co-expression summary of targets of transcription factors TF_z1_ and TF_z2_, which interact with TF_x_.

**Supplementary Table 3.** Co-expression summary of targets of transcription factors TF_z1_ and TF_z2_, which interact with TF_x_ and with each other

**Supplementary Table 4.** Total number of genes highly co-expressed (HCG) with TF_x_ that are also targets of TF_z_.

## Notes

### Competing Interest Statement

The authors have declared no competing interest.

